# Characterization of SID-1-dependent and independent intergenerational RNA transport pathways in *Caenorhabditis elegans*

**DOI:** 10.1101/156232

**Authors:** Eddie Wang, Craig P. Hunter

**Affiliations:** Department of Molecular and Cellular Biology, Harvard University, Cambridge, MA 02138

**Keywords:** *Caenorhabditis elegans*, RNAi, double-stranded RNA, yolk

## Abstract

Systemic RNA interference (RNAi) in *C. elegans* is dependent on *sid-1* (Winston *et al.* 2002), *sid-3* (Jose *et al.* 2012) and *sid-5* (Hinas *et al.* 2012). After injection, expression, or ingestion, double-stranded RNA (dsRNA) is transported between cells throughout the animal to enable RNAi in most tissues, including the germline and progeny. Here, we characterize the role of the Sid genes in transport of dsRNA to progeny. We previously reported that dsRNA injected directly in the germline unexpectedly requires *sid-1* activity in the progeny to initiate RNAi (Winston *et al.* 2002). We now show that germline injected dsRNA can travel by three independent pathways to silence gene expression in embryos. First, germline injected dsRNA is delivered, presumably by bulk flow, into oocytes and embryos. This means of delivery, which does not require *sid-1,* is limited by the amount and location of injected dsRNA. Second, maternal *sid-1* transports extracellular dsRNA into the germline where it can silence maternal deposited mRNAs and segregate to embryos to silence embryonically expressed mRNAs. Third, extracellular dsRNA is also endocytosed into oocytes by the low-density lipoprotein (LDL) receptor superfamily homolog RME-2. The endocytosed dsRNA then requires *sid-1* and *sid-5* in embryos to silence embryonically expressed genes. Extracellular fluorescent dsRNA, once endocytosed into oocytes, does not co-localize with VIT2::GFP and it does not require *sid-1* activity to segregate from the late endocytosis marker GFP::RAB-7 in early embryos. In conclusion, we identify genes and pathways that function redundantly for intergenerational RNA transfer that may represent mechanisms for transgenerational epigenetic inheritance.

## Introduction

RNA interference is a powerful and well-conserved mechanism for sequence specific gene silencing (HutvÁgner and Zamore 2002). Introduced double stranded RNA (dsRNA) triggers degradation of homologous transcripts (Fire *et al.* 1998; Hamilton and Baulcombe 1999; Zamore *et al.* 2000; Sharp 2001) as well as subsequent transcriptional gene silencing (Guang *et al.* 2008; Guang *et al.* 2010; Buckley *et al.* 2012). In some animals including the nematode *C. elegans*, RNAi is systemic; dsRNA introduced into the animal by any of several methods results in rapid spread of silencing throughout the animal (Fire *et al.* 1998; Ivashuta *et al.* 2015). This systemic silencing requires the dsRNA channel SID-1, which imports dsRNA into the cytoplasm (Winston *et al.* 2002; Feinberg and Hunter 2003; Shih and Hunter 2011). SID-1 supports particularly effective silencing in the progeny of dsRNA exposed mothers (Fire *et al.* 1998; Grishok *et al.* 2000; Alcazar *et al.* 2008), implying transfer of dsRNA from mother to embryo.

SID-1 dependent heritable RNAi can be initiated by ingesting dsRNA expressing bacteria, by tissue-specific expression of dsRNA from transgenes, and by dsRNA injection into somatic tissues, the germline, or the pseudocoelom (PC, body cavity) (Jose *et al.* 2009). Initial investigation of *sid-1*-dependent heritable RNAi by dsRNA injection showed that dsRNA targeting a germline expressed mRNA injected into the gonad of *sid-1* mutant animals produced affected progeny, while injection into the intestine or the PC produced no affected progeny (Winston 2002; Winston *et al.* 2002). Similar injections into wild-type animals produced near 100% affected progeny. These results showed that *sid-1* is required for transport of dsRNA into the germline, but is not required for RNAi itself in the germline. Similar results were expected for injection of dsRNA targeting the somatic muscle gene *unc-22*, which is required for normal muscle function. However, while *unc-22* dsRNA injection into wild-type animals produced many affected progeny, similar injections into *sid-1* mutants failed to produce affected progeny, even among the progeny of germline injected mothers. This was unexpected because the syncytial germline cytoplasm is continuous with the oocyte/embryo cytoplasm and thus SID-1 should not be required. Also unexpectedly, crossing wild-type males to *unc-22* dsRNA injected *sid-1* mutant hermaphrodites restored silencing to the progeny. This was true for both germline injected mothers (Winston *et al.* 2002) as well as PC injected mothers (Winston 2002). These results revealed a role for *sid-1* in the embryo and an unidentified *sid-1*-independent pathway for dsRNA transport to embryos.

Here we report a detailed genetic and cytological investigation of dsRNA transport into oocytes and embryos. We identify three dsRNA transport process that support inherited RNAi. First, as expected, germline injected dsRNA segregates, independent of *sid-1* activity, to embryos resulting in temporally limited silencing. Second, maternally expressed *sid-1* transports extracellular dsRNA into the germline. Third, the LDL receptor superfamily homolog RME-2 enables endocytosis of dsRNA into oocytes, but to initiate RNAi in the resulting embryos zygotic *sid-1* and *sid-5* are required, presumably to release membrane encapsulated dsRNA into the cytosol. Marré *et al.* recently confirmed the published observations (Winston 2002; Winston *et al.* 2002) on an embryonic role for *sid-1* and the presence of a *sid-1*-independent transport process and also identified RME-2 as important for this *sid-1* independent dsRNA transport (MarrÉ *et al.* 2016). In contrast to Marré *et al.*, our analysis shows that maternal RME-2 and SID-1 act independently, as neither single mutant prevents dsRNA transport to embryos. Our analysis of this discrepancy revealed a strong effect of maternal developmental stage on inherited RNAi, which only strengthens the discrepancy. Our analysis of injected labeled dsRNA shows that although dsRNA and the yolk marker VIT-2::GFP co-localize in the PC space and even on the surface of the oocyte, internalized VIT-2::GFP and dsRNA do not co-localize. Furthermore, labeling dsRNA with Cy5 interferes with dsRNA transport into oocytes. This indicates that non-specific interactions between yolk and dsRNA are unlikely to account for the RME-2 mediated uptake. Our genetic analysis of post-endocytosis dsRNA trafficking shows that dsRNA either exits the endocytosis pathway early or rapidly transits the pathway independently of *sid-1*, as dsRNA and late endosome markers do not significantly co-localize in either wild-type or *sid-1* mutant embryos.

## Results

### Inherited silencing in the absence of *sid-1*

SID-1 is required to transport dsRNA or derived silencing signals to the germline, as shown by injecting dsRNA into specific tissues or the PC (Winston 2002; Winston *et al.* 2002). However, while *unc-22* dsRNA injection into any tissue in wild-type animals resulted in silenced progeny, *unc-22* dsRNA injected into the syncytial germline of a *sid-1* mutant, unexpectedly, did not result in any affected progeny (Winston 2002; Winston *et al.* 2002). To further investigate this result, we injected *unc-22* dsRNA directly into the syncytial gonads of wild-type and *sid-1* mutant adult hermaphrodites and every two hours after injection collected their self-progeny. As expected, and consistent with previous observations of systemic silencing, the proportion of twitching progeny from injected wild-type animals quickly rose to 100% and was sustained for the duration of the experiment (Fig. 1*B*). In contrast to previous observations, we found that injecting dsRNA directly into the syncytial gonads of *sid-1^-/-^* animals produced strongly twitching progeny. However, this was only true of embryos laid within approximately the first 18 hours after injection (Fig. 1*A*). The timing of peak silencing varied between injected P_0_ animals, but always reached 100% (Fig. *S*1). The subsequent decrease in fraction of silenced F_1_ embryos to zero suggests that the injected *unc-22* dsRNA is rapidly depleted. In the previous experiments (Winston 2002; Winston *et al.* 2002), the injected hermaphrodites were allowed to recover for up to 24 hours before progeny were collected and scored for silencing. The apparent rapid depletion of syncytial germline injected dsRNA explains the past failure to detect *sid-1*-independent silencing. Furthermore, when a single gonad arm was injected in *sid-1* mutant animals, only a maximum of 50% twitching progeny was produced (Fig. 1*C*, *S*1). Thus, *sid-1* independent silencing is restricted to the cytoplasm containing the injected dsRNA. In contrast, the control injections into a single gonad of wild-type hermaphrodites clearly show gonad injected dsRNA is mobile between gonad arms (Fig. 1*D*). Additionally, in wild type animals, dsRNA or a derived silencing signal persists indefinitely in the injected hermaphrodites. Interestingly, injecting *unc-22* dsRNA into a single gonad arm of *sid-1* mutant hermaphrodites and then crossing them to wild-type males produced nearly 100% affected heterozygous cross progeny (Fig 1*E*). This indicates that *sid-1* activity in the mother is not required for either gonad to gonad transfer of dsRNA or the long-term persistence of the silencing signal, and that *sid-1* activity in the embryo is required to access this dsRNA or derived silencing signal.

**Fig. 1.**
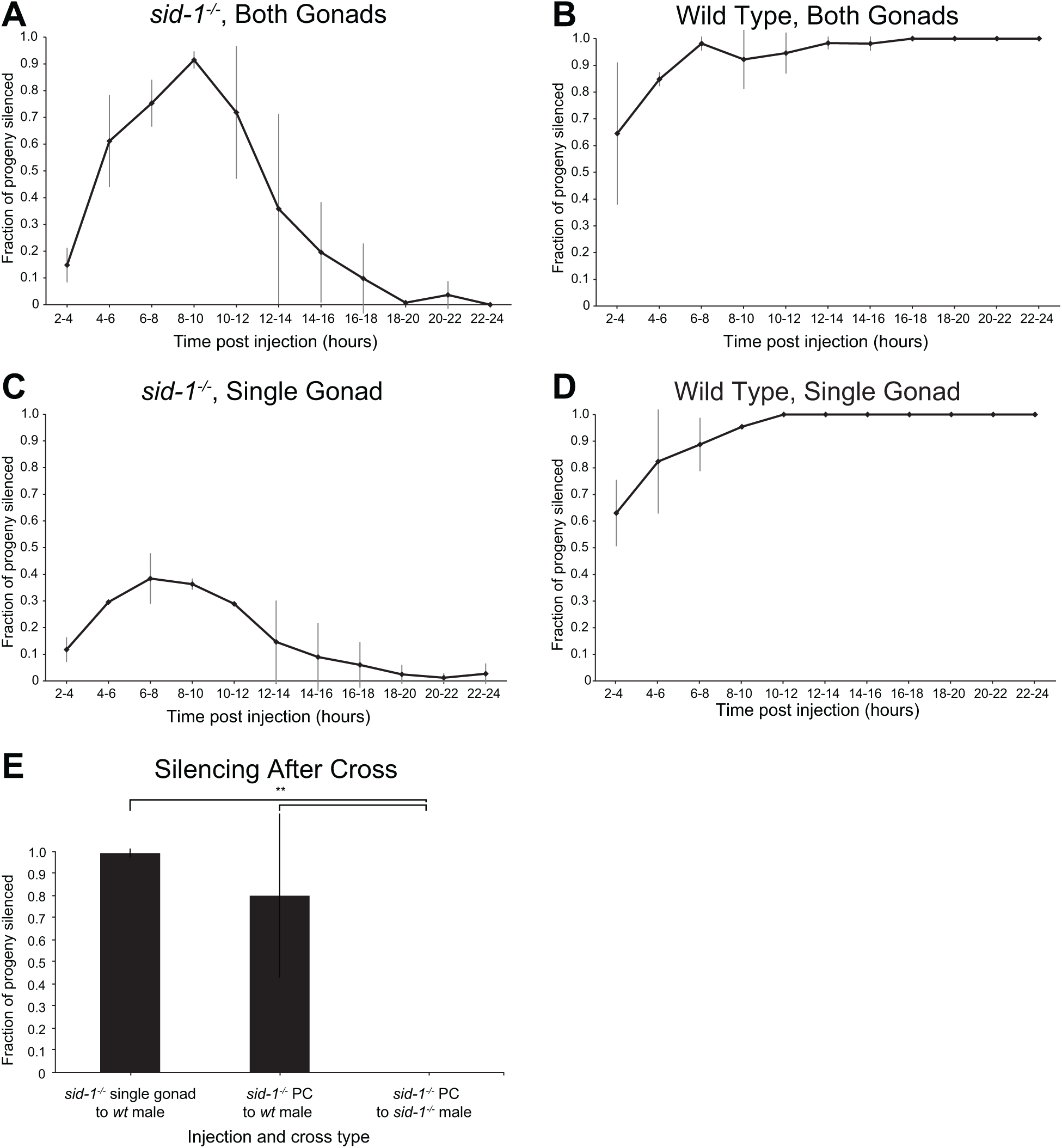
*sid-1*-dependent and -independent silencing in progeny of dsRNA injected parents. (*A-D*) Time course of fraction of progeny with the Unc-22 phenotype laid after *unc-22* dsRNA gonad injection into wild-type or *sid-1* mutant hermaphrodites. (*E*) Fraction of progeny with Unc-22 phenotype following *unc-22* dsRNA gonad or PC injection into *sid-1^-/-^* mutant hermaphrodites crossed to wild-type or *sid-1* mutant males. Error bars in (*A-D*) represent standard error from two experiments with 10 injected hermaphrodites each. Error bars in (*E*) represent standard deviation from 4, 6, and 3 injected hermaphrodites, left to right. \*\**p < 0.01* by Welch’s *t* test.

### *sid-1*-independent dsRNA Transmission Requires RME-2

The above results, showing *sid-1*-independent delivery of dsRNA from mothers to embryos and a *sid-1*-dependent step in the embryos for effective RNAi is similar to feeding RNAi, where the uptake and transport of ingested dsRNA is separable. Feeding RNAi first requires SID-2, an intestinal-lumen-localized transmembrane protein, for endocytosis of ingested dsRNA, and SID-1 is required subsequently for effective RNAi within intestinal cells, likely to release dsRNA from endosomes (Winston *et al.* 2007; Mcewan *et al.* 2012). A candidate receptor for similar dsRNA endocytosis in the germline is RME-2, an LDL receptor superfamily homolog that functions in oocytes as a yolk/lipoprotein receptor. In *C. elegans,* yolk is synthesized in the intestine, exported to the PC, and then taken up by oocytes via receptor-mediated endocytosis (Sharrock *et al.* 1990; Grant and Hirsh 1999). Thus, we hypothesized that PC dsRNA may bind to either yolk protein, another RME-2 substrate, or directly to RME-2 for endocytosis into oocytes. To test this hypothesis, we examined *rme-2^-/-^* mutants, which do not take up any yolk (Grant and Hirsh 1999) We injected *unc-22* dsRNA into the PC of *rme-2^-^ sid-1^-/-^* double mutant hermaphrodites and then crossed these animals to wild-type males. The heterozygous cross progeny did not twitch, showing that RME-2 is required for heritable RNAi (Fig. 3A). This indicates that RME-2 can endocytose PC dsRNA into oocytes. Similar findings were recently reported by Marré *et al.* (MarrÉ *et al.* 2016).

### SID-1 is sufficient in the germline to transmit maternal dsRNA to progeny

SID-1 and SID-2 are both individually required for feeding RNAi (Winston *et al.* 2007). That is, SID-1 is not sufficient to transport ingested dsRNA into intestinal cells. To determine whether RME-2 and SID-1 must similarly function together to deliver PC dsRNA to the silencing components in embryos we injected *unc-22* dsRNA into the PC of *rme-2^-/-^* single mutants. We found that their progeny showed Unc-22 defects (Fig. 3*A*). This result indicates that, unlike the case in the intestine, SID-1 may directly transport PC dsRNA to oocytes and embryos.

This finding contrast with recent results reporting that *rme-2* single mutants were defective for inherited RNAi initiated by consuming *unc-22* dsRNA (MarrÉ *et al.* 2016). The difference in potency between feeding RNAi used by Marré *et al.,* and PC injection of concentrated *in vitro* transcribed dsRNA as described above may explain their negative result. To address this, we attempted to repeat their feeding RNAi assay as described, *i.e.* exposing L4 hermaphrodites to *unc-22* dsRNA-expressing bacteria for one day before washing and transferring to control bacteria. By these methods, we failed to detect robust inherited RNAi among the progeny of wild-type parents. Further analysis revealed a striking dependency on maternal developmental stage for inherited silencing. We split a batch of freshly hatched wild-type larvae into 3 populations and exposed each to *unc-*22 RNAi food on either day 1 (L1/L2), day 2 (L3/L4), or day 3 (adult) after hatching, thoroughly washing off residual bacteria each day. After a final wash, animals were allowed to lay F_1_ progeny on non-RNAi food. We found that only animals exposed to *unc-22* food as adults efficiently produced twitching progeny (Fig. 3B), even though the P_0_ animals fed on days 1 and 2 still exhibited a strong Unc-22 twitching phenotype immediately after RNAi exposure. Using these conditions, we found that the progeny of wild type and *rme-2* adults placed on *unc-22* food show similar proportions of twitching progeny (Fig. 3C). Similarly, *rme-2* L4 animals exposed to RNAi do not produce twitching progeny, in agreement with Marré *et al*. Because inherited RNAi by dsRNA feeding requires adult exposure, it is reasonable to assume that the slightly slower development of *rme-2* mutants differentially limited the period of adult exposure compared to wild-type animals when initially placed on the RNAi food as L4 larvae. In conclusion, independent of the means of dsRNA delivery, *rme-2* is not required for transport of dsRNA into the germline.

### *sid-5* functions with *sid-1* in the embryo

Pseudocoelomic *unc-22* dsRNA transported into oocytes or embryos in the absence of maternal *sid-1* requires embryonic *sid-1* to silence *unc-22*. We next asked whether *sid-2*, *sid-3*, or *sid-5* are also required in the embryo. We first crossed *sid-1; sid-2* double mutant hermaphrodites injected with *unc-22* dsRNA to either wild type or *sid-2* males and scored silencing in the cross progeny. If *sid-2* activity is required in the embryo, then the cross progeny of only the wild type males will twitch. If *sid-2* activity is not required in the embryo, then the cross progeny of both wild type and *sid-2* males will twitch. Using this method we determined that neither *sid-2* nor *sid-3* are required for either maternal uptake or embryonic release (Fig. 2A,B), but we found a striking requirement for *sid-5* for embryonic release.

**Fig. 2.**
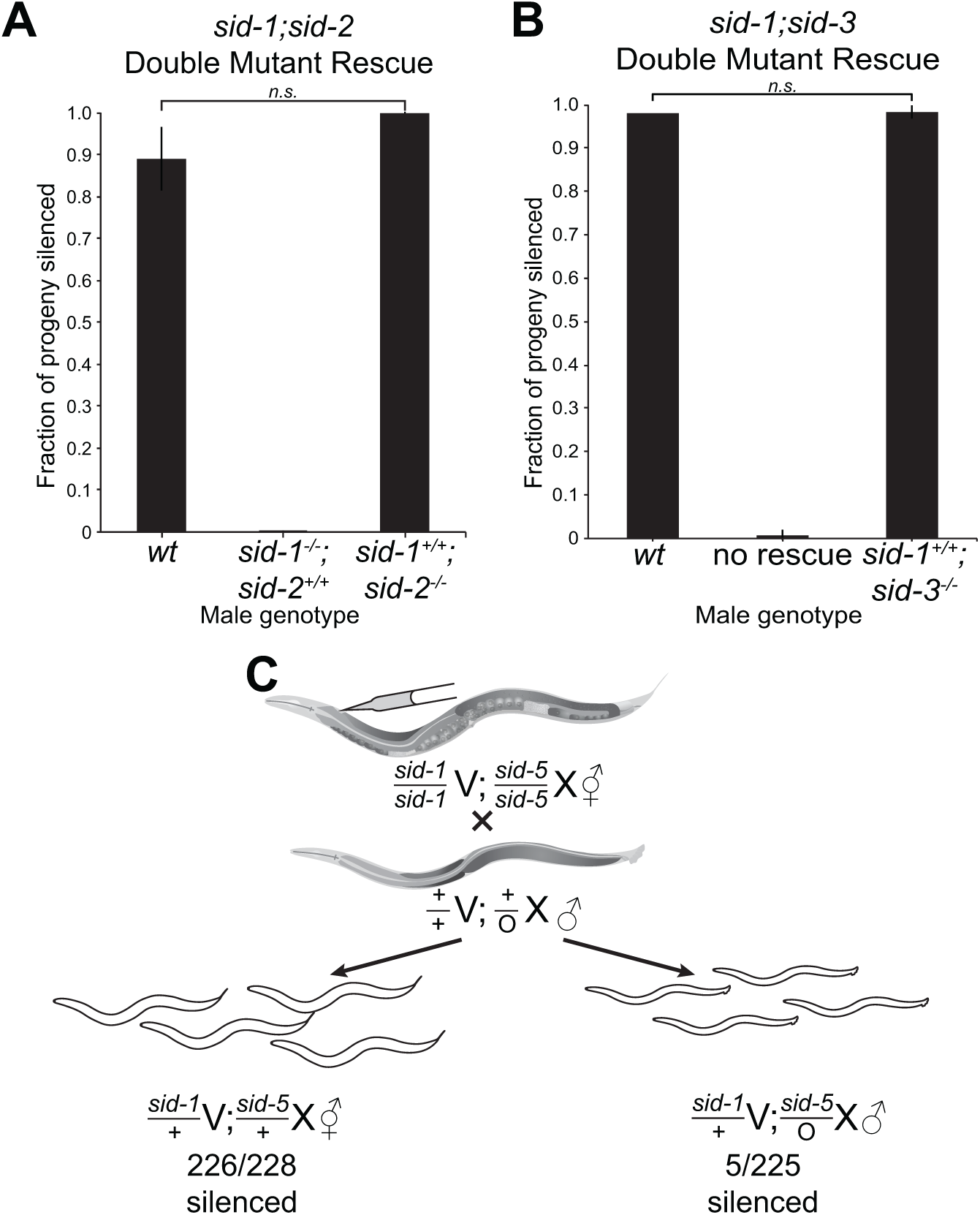
Maternal and zygotic Sid-dependence of inherited silencing. (*A*) Fraction of *unc-22* silenced cross progeny from *sid-1; sid-2* double-mutant hermaphrodites first PC-injected with *unc-22* dsRNA and then crossed to either wild-type, *sid-1; sid-2* double mutant, or *sid-2* single mutant males. (*B*) Fraction of *unc-22* silenced cross progeny from *sid-1; sid-3* double-mutant hermaphrodites first PC-injected with *unc-22* dsRNA and then crossed to wild-type, *sid-1; sid-3* double mutant, or *sid-3* single mutant males. (*C*) Schematic of the injection, cross, and *unc-22* silencing scoring of cross progeny from *sid-1; sid-5* double mutant hermaphrodites first PC-injected with *unc-22* dsRNA and then crossed to wild-type males. *sid-5* is X-linked, thus hermaphrodite progeny are heterozygous and males are hemizygous. Error bars in (*A, B*) represent standard deviation from 4, 7, and 5 injected hermaphrodites in (*A*) and 1, 3, and 3 injected hermaphrodites in (*B*). 3 injected hermaphrodites in (*C*). *n.s.* = not significant.

**Fig. 3.**
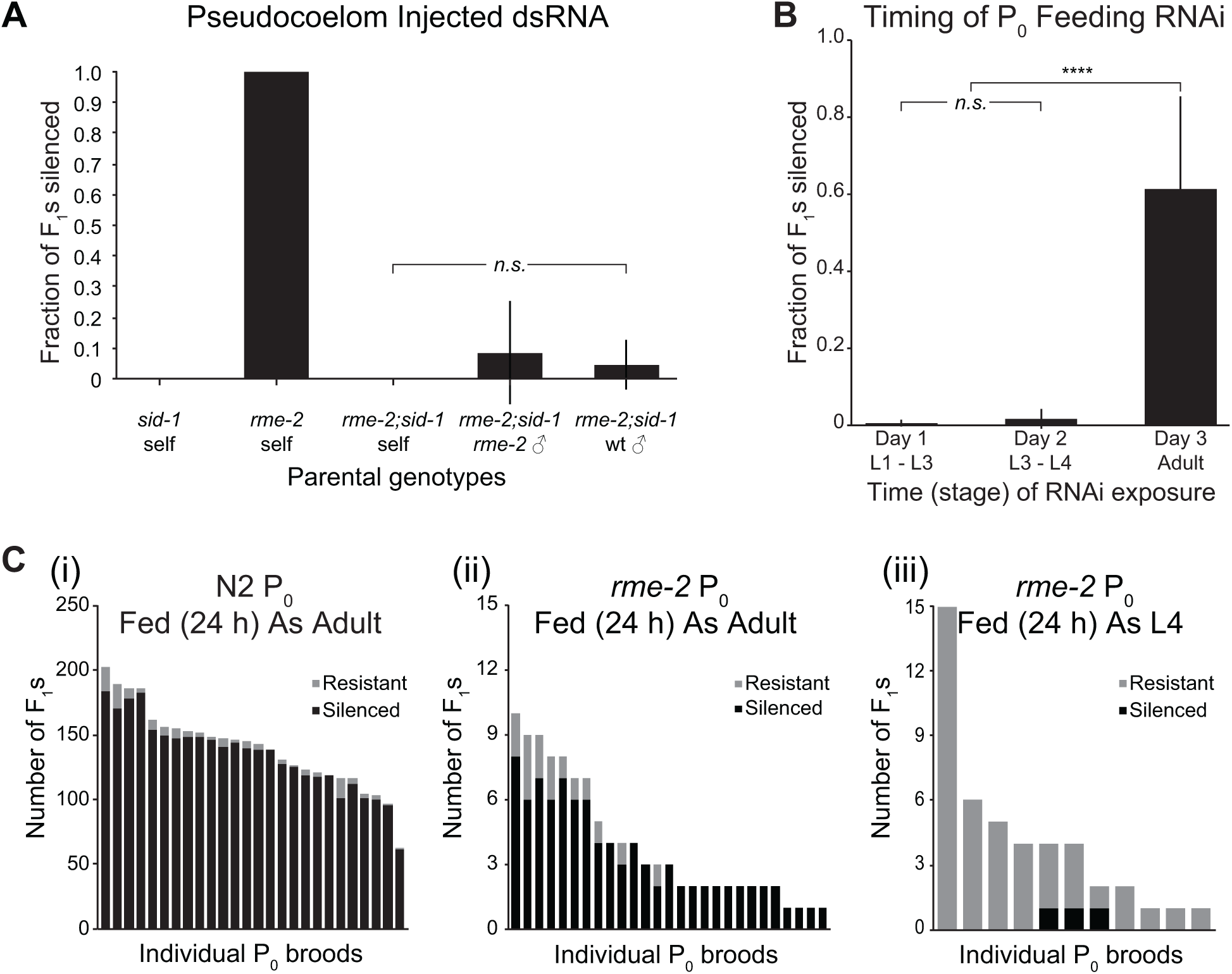
Maternal RME-2-dependent inherited silencing. (*A*) Fraction of progeny sensitive to *unc-22* silencing among the self-progeny or indicated cross-progeny of hermaphrodites PC-injected with *unc-*22 dsRNA. *n* = 6, 5, 3, 4, and 12 injected hermaphrodites respectively. (*B*) Fraction of progeny sensitive to *unc-22* silencing after wild-type parents were exposed to feeding RNAi at the given periods of time after hatching. *n* = 30 treated parents for each group. (*C*) Sensitivity to *unc-22* feeding RNAi in progeny after treating (*i*) wild-type parents as adults, (*ii*) *rme-2* mutant parents as adults, or (*iii*) *rme-2* mutant parents as L4 larvae. Because *rme-2* mutants have severely reduced fecundity, the results from each individual parent are presented separately for clarity, with silenced progeny represented in black bars and non-silenced progeny in grey bars. All error bars represent standard deviation. \*\*\*\**p* < *0.00001* by *t* test. *n.s.* = not significant.

SID-5 is an endosome-associated protein required for efficient systemic RNAi (Hinas *et al.* 2012). In our double mutant rescue experiments, no silencing was observed in the next generation if *sid-5* is not rescued (Fig. 2*C*), the same effect as if *sid-1* is not rescued. Because *sid-5* is located on the X-chromosome, crossing a *sid-1 ^-/-^; sid-5^-/-^* hermaphrodite to a wild-type male results in heterozygous *sid-5^+/-^* hermaphrodite progeny and hemizygous *sid-5^0/-^* mutant male progeny. When performing the *unc-22* dsRNA PC injection and rescue in this context, we saw that 100% of hermaphrodite progeny but none of the male progeny showed the *unc-22* silencing phenotype (Fig. 2*C*). The importance of SID-5 in the context of *sid-1*-independent transmission of RNA is especially surprising given the previous reports of only weak systemic RNAi defects for *sid-5* mutants (Hinas *et al.* 2012). However, those experiments were performed in the context of fully functional and normally expressed SID-1, in which case there may be alternative routes for RNA to reach the cytoplasm that are less dependent on SID-5.

### Labeled Nucleotides Can Be Used to Visualize Functional Transported dsRNA

Our finding that RME-2 is required for silencing signals to reach developing oocytes and embryos suggests an association between RNA and yolk. We sought evidence of this association by labeling and visualizing both components. We synthesized *unc-22* dsRNA using 5-ethynyluridine (5EU) nucleotides, replacing a significant fraction of the normal uridine nucleotides with nucleotides carrying the modified base. 5EU carries a small alkyne modification that allows the RNA to be easily visualized after fixation and labeling through click chemistry (Jao and Salic 2008). 5EU-labeled *unc-22* dsRNA injected into a single gonad arm of a wild-type worm results in >50% twitching progeny, indicating that, like unmodified dsRNAs, 5EU dsRNA is both mobile and capable of silencing target genes (Fig 4*A*).

**Fig. 4.**
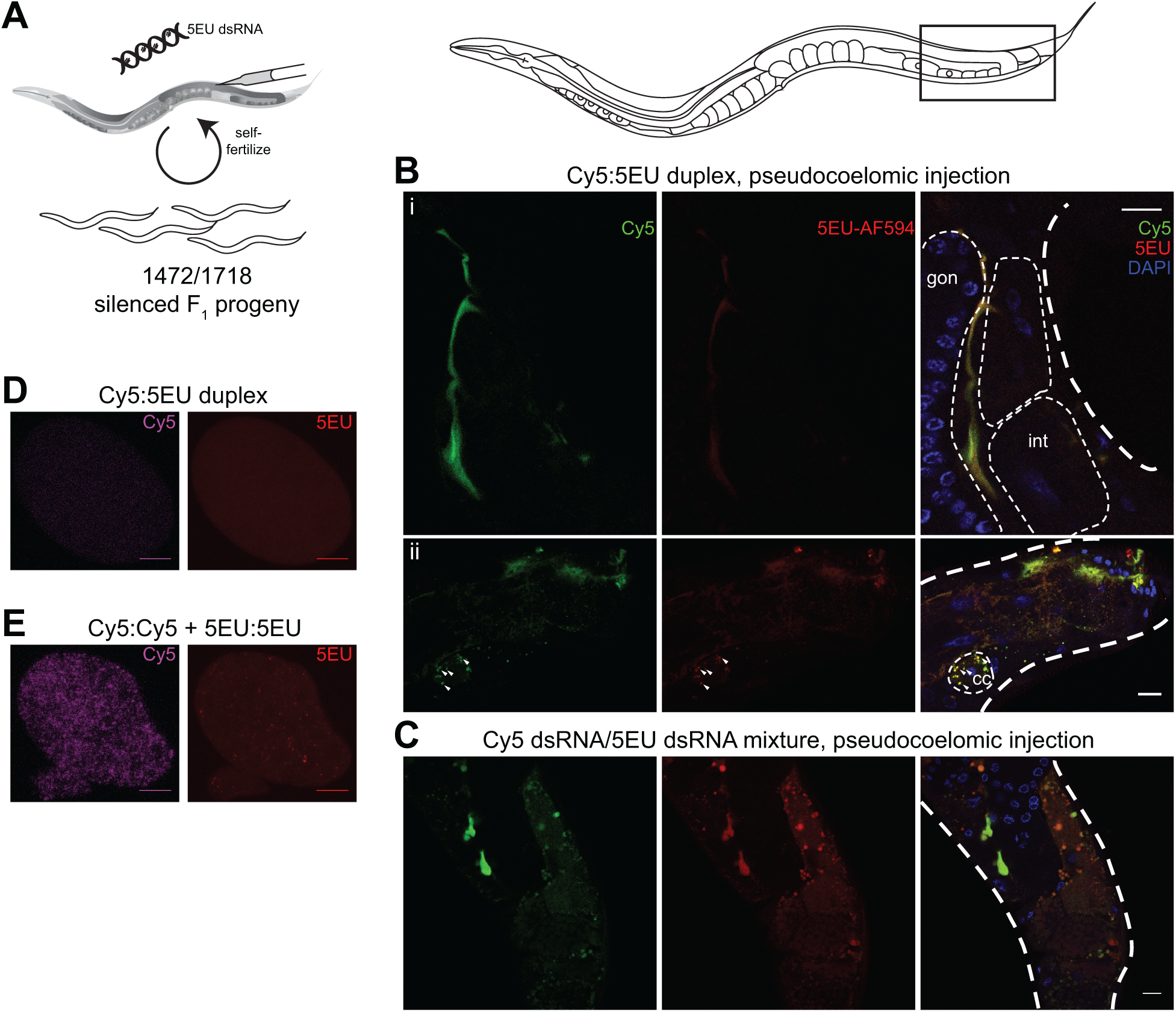
Visualizing 5-Ethynyluridine (5EU) labeled functional dsRNA. (*A*) Injected 5EU dsRNA injected into only one gonad arm produces >50% affected progeny. *n* = 8 injected hermaphrodites. (*B*) Localization of PC injected Cy5::5EU heteroduplex dsRNA. Cy5 fluorescence and 5EU detection co-localize in the pseudocoelom (*i*) and a coelomocyte (*cc)* (ii; white arroweads). (*C*) Independent localization of PC injected 5EU and Cy5 labeled dsRNA. Images in (*B*) and (*C*) represent portions of dissected and partially flattened adult hermaphrodites. Thick dotted lines mark the boundary of the animal, and thinner dotted lines mark structures such as the gonad (*gon*) or intestinal cells (*int*) as landmarks for orientation. (*D, E*) Cy5 and 5EU signal in embryos collected from adults injected with the dsRNA species described in (*B*) and (*C*) respectively. The two Cy5 images are overexposed, revealing diffuse autofluorescence and no detectable RNA. Scale bars = 10 μm.

To determine whether fluorescently labeled 5EU signal represents intact dsRNA and not degraded nucleotides or other irrelevant species, we used a second label and synthesized dsRNA composed of one strand labeled with 5EU and the other strand internally labeled with Cy5-uridine. Although Cy5-labeled dsRNA is not capable of systemic RNAi, it suffices for demonstrating the properties of labeled dsRNA. We injected the Cy5:5EU duplex dsRNA into the PC of adult hermaphrodites and several hours later fixed the animals to visualize both labels (Fig. 4*B*). Much of the signal appears in the pseudocoelom as pools surrounding other tissues. Importantly, both labels are detected in the same locations and their intensities appear closely correlated. Coinciding labeled RNA was also detected in punctate foci in coelomocytes, providing further evidence for the structure and stability of the injected RNA.

In addition to the Cy5:5EU duplex, we also injected a 1:1 mixture of Cy5-labeled dsRNA with 5EU-labeled dsRNA. We found that although these mixed dsRNAs were both readily detectable in the PC, their apparent relative abundances were much less well-correlated than the co-labeled duplex dsRNA, and more frequently one label could be seen without the other (Fig. 4*C*). That is, when not physically bound together, each label appears free to vary independently. Together, these data suggest that 5EU fluorescence represents *bona fide* dsRNA as opposed to disassociated strands or degraded nucleotides or labels.

We also attempted to detect labeled RNA in embryos from PC-injected hermaphrodites. The Cy5:5EU heteroduplex was not detected above background in embryos (Fig. 4*D*). Furthermore, while we could detect 5EU-labeled RNA punctae in embryos, the co-injected Cy5-labeled RNA was only detected in the PC and coelomocytes (Fig. 4*E*). These observations are consistent with Cy5-containing RNA being impaired for normal trafficking while 5EU-labeled RNA behaves as expected for unmodified RNA. The inability of either the Cy5:5EU heteroduplex or Cy5-RNA to enter embryos suggests there is selectivity in the RME-2-dependent transmission process. Such selectivity is not expected if dsRNA is simply hitchhiking on yolk proteins.

### Labeled dsRNA and VIT-2::GFP largely fail to co-localize

We next sought to characterize the relationship between dsRNA and yolk. Previous studies have shown co-localization of yolk and end-labeled dsRNA that lessens after fertilization (MarrÉ *et al.* 2016). We injected our 5EU-labeled dsRNA into the PC of adults expressing GFP-labeled yolk (*vit-2::GFP*) and imaged the injected adults as well as isolated embryos. We detected 5EU-labeled dsRNA co-localized with VIT-2::GFP granules at the surface of developing oocytes in adult gonads (Fig. 5*A*), in agreement with prior observations (MarrÉ *et al.* 2016). This co-localization is consistent with co-dependence on oocyte-expressed RME-2 for the uptake of yolk and RNA. However, interior sections contain 5EU punctae not associated with any yolk, suggesting that although they may share the same endocytosis receptor, they are either separately endocytosed or sorted to different endosomes soon after import. While previous studies reported loss of co-localization in embryos past the 4-cell stage (MarrÉ *et al.* 2016), higher resolution imaging suggests that the separation of yolk and dsRNA can be seen even earlier in oocytes soon after import (Fig. 5*A*). Analysis of 5EU-labeled dsRNA and VIT-2::GFP in embryos is consistent with this view. The majority of detectable 5EU fluorescence in embryos does not co-localize with GFP fluorescence (Fig. 5*B*). The lack of co-localization might reflect *sid-1*-dependent release of dsRNA from endosomes. However, in *sid-1^-/-^* mutant embryos labeled dsRNA also fails to co-localize with VIT2-GFP (Fig. 5*B*). This further supports the idea that although dsRNA and yolk share a common mechanism for endocytosis into oocytes, they likely do not share common endosomal compartments.

**Fig. 5.**
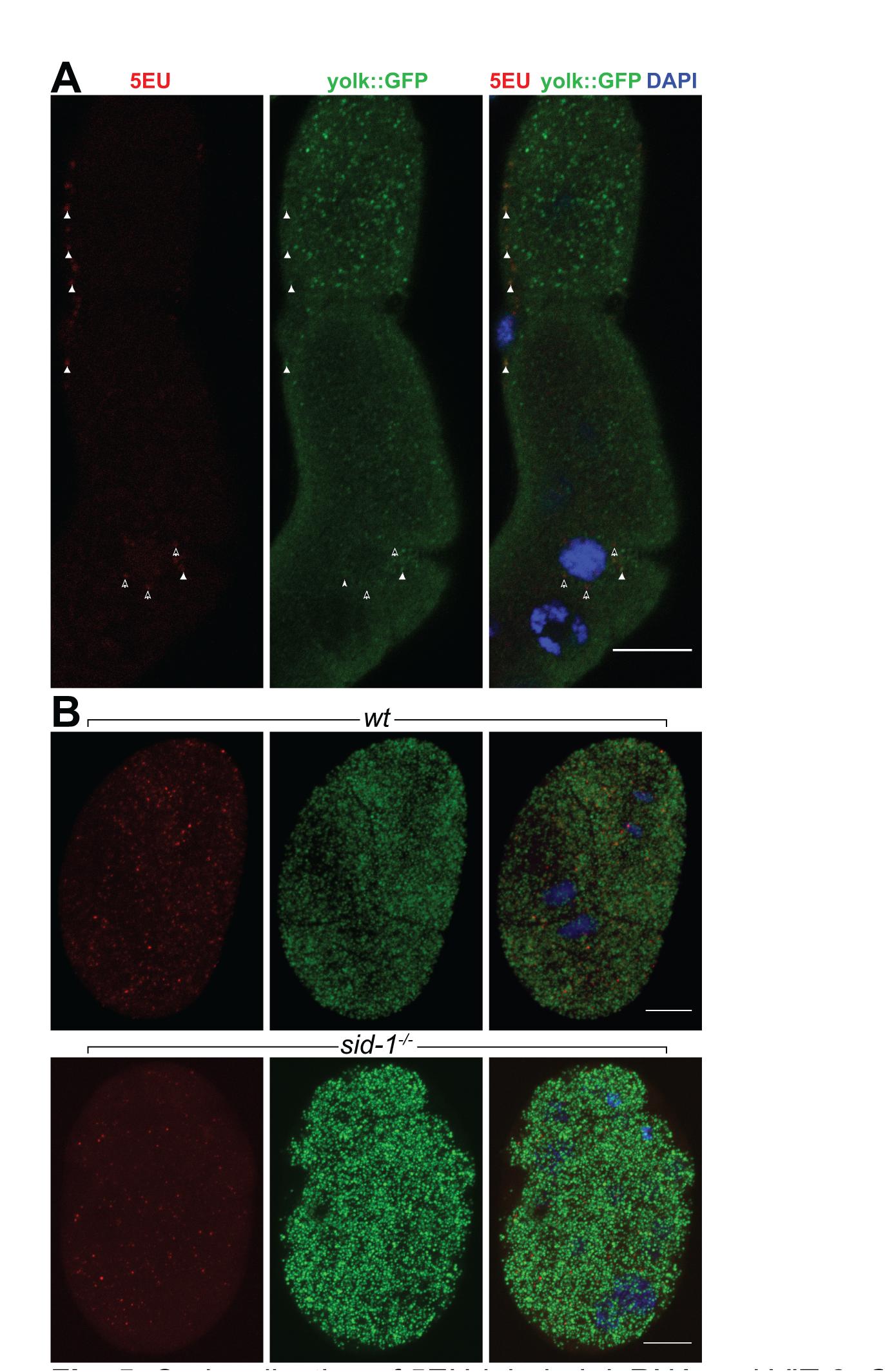
Co-localization of 5EU-labeled dsRNA and VIT-2::GFP on and within oocytes. (*A*) PC-injected 5EU-labeled dsRNA co-localized with GFP-labeled yolk at the surface of developing oocytes (white arrowheads), but not intracellularly (notched arrowheads). More proximal oocyte (top) contains more VIT-2::GFP. (*B*) Maximum *z*-projections of VIT-2::GFP and 5EU-labeled dsRNA in wild-type and *sid-1^-/-^* embryos show little co-localization. Scale bars = 10 μm.

### dsRNA Transits RAB-7-Containing Vesicles Independent of SID-1

Since internalized dsRNA appears to be physically separate from yolk granules, we wondered whether the 5EU punctae might co-localize with the late endosome marker RAB-7 (Feng *et al.* 1995; Grant and Hirsh 1999). We injected adult hermaphrodites in the PC with 5EU-labeled dsRNA and visualized the resulting embryos in conjunction with antibody staining against GFP-labeled RAB-7. In wild-type embryos, we occasionally detected dsRNA together with RAB-7-positive vesicles, but the majority of the 5EU punctae were not associated with GFP-RAB-7 (Fig. 6*A*, upper row). In *sid-1* mutant embryos, 5EU punctae are also readily detected outside of RAB-7 vesicles, and there does not appear to be significantly more 5EU retained within RAB-7 vesicles (Fig. 6*A*, lower row). Thus, although silencing by inherited dsRNA involves early steps in the endocytotic pathway, the dsRNA appears to transit this system rapidly. Although *sid-1* is required for inherited silencing, its activity is not apparent in the localization of 5EU-labeled dsRNA in oocytes or early embryos.

**Fig. 6.**
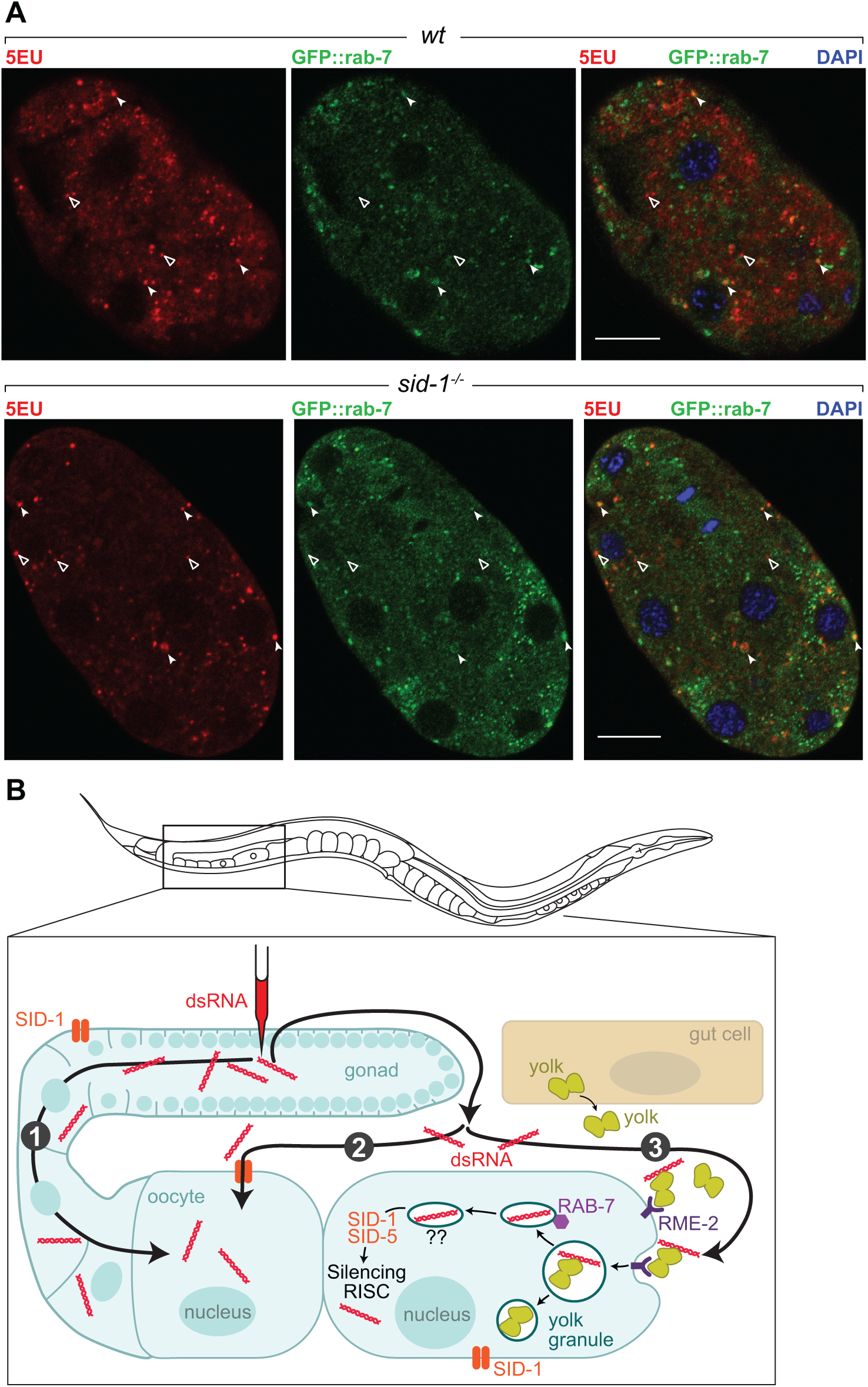
Co-localization of dsRNA and GFP::RAB-7. (*A*) 5EU foci detected in embryos after pseudocoelomic 5EU-labeled dsRNA injection co-localizes with GFP:: RAB-7 (notched arrowheads) but is also found outside of RAB-7 structures (open arrowheads) in both wild-type (upper row) and *sid-1* mutant (*Lower row*) embryos. Scale bars = 10 μm. (*B*) Model for three inherited dsRNA transport pathways. 1) DsRNA injected directly into the syncytial germline can silence the resulting progeny without SID-1. Some injected dsRNA exits the gonad to the PC and is then 2) endocytosed into developing oocytes via LDL receptor super-family member RME-2, or 3) this PC dsRNA can also be directly transported into oocytes via SID-1.

## Discussion

Intercellular dsRNA transport in *C. elegans* via the dsRNA channel SID-1 supports systemic gene silencing throughout the animal and its progeny. Here, we have described and characterized the maternal and embryonic processes that support intergenerational dsRNA transport, as summarized in figure 6*B*.

First, we more fully described the result of dsRNA injection into the gonad. In wild-type animals, dsRNA injection into a single gonad arm results in robust silencing in nearly all progeny, while similar injection in *sid-1* mutant results in transitory silencing and only in progeny derived from the injected gonad arm (Figure 1; (Winston 2002; Winston *et al.* 2002). While we still do not understand how injected dsRNA escapes the injected gonad arm or its long-term persistence within the injected animals, its subsequent uptake into both gonad arms can occur by either a *sid-1*-dependent process or a *sid-1*-independent but *rme-2*-dependent process (Figure 2). Consistent with the role of RME-2 as an endocytosis receptor, *sid-1* activity is required subsequently, presumably to release dsRNA from a membrane enclosed compartment (Figure 3). A recent report concluded that silencing in progeny in response to ingested dsRNA is dependent on *rme-2* (MarrÉ *et al.* 2016). Because this conflicts with our results, we repeated their analysis, discovering that progeny silencing in response to parental feeding RNAi is developmentally restricted, being most effective in adults. We found that the progeny of *rme-2* adults are efficiently silenced by feeding RNAi, and suggest that the slowed development of *rme-2* animals may have compromised their analysis (Figure 3).

The redundancy between *sid-1* and *rme-2* for uptake and transport of dsRNAs into the germline is contrasted by the necessity for both *sid-1* and *sid-2* for uptake and transport of dsRNAs into the intestine (Winston *et al.* 2007). Presumably, oocyte expressed SID-1 localizes to the plasma membrane adjacent to the PC space, whereas intestinal expressed SID-1 does not localize the lumenal plasma membrane. This difference may reflect the privileged environment of the PC space, relative to the ingested intestinal milieu. An alternative but unlikely scenario is that a second oocyte expressed endocytosis receptor accounts for the *rme-2*-independent uptake. However such activity should be apparent in silencing the progeny of *rme-2; sid-1* double mutants, which is not detected. The redundancy between *sid-1* and *rme-2* is significant in light of reports of *sid-1*-independent RNAi-dependent transgenerational inheritance (Schott *et al.* 2014). It will be interesting to determine whether alternative RNA uptake mechanisms mediate transgenerational inheritance and inheritance of acquired traits.

Our genetic and cytological analysis of dsRNA in oocytes and embryos revealed a role for *sid-5* alongside *sid-1* in dsRNA import and showed that despite sharing an endocytosis receptor, yolk protein and dsRNA travel independent intracellular routes. SID-5 is a late endosomal/multivesicular body localized protein required for efficient systemic RNAi (Hinas *et al.* 2012). Tissue-specific rescue experiments showed that *sid5* expression in the intestine but not the muscle was sufficient for RNAi silencing of a muscle gene (Hinas *et al.* 2012). The results presented here, showing a role for SID-5 in dsRNA import, suggest that efficient release of endocytosed ingested dsRNA to the cytoplasm is important for subsequent export to muscle cells. Our cytological analysis of dsRNA within oocytes and embryos indicates that yolk and dsRNA rarely co-localize (Figure 5). This result challenges the assumption that dsRNA enters ooctyes with yolk granules. Furthermore, the observation that EU- but not Cy5-labeled dsRNA injected into the PC is internalized within oocytes (Figure 4) suggest specificity in dsRNA uptake rather than proposed non-specific mechanisms (MarrÉ *et al.* 2016). While *sid-1* and *sid-5* are required for this endocytosed dsRNA to trigger RNAi silencing, *sid-1* activity is apparently not required for transit to post-RAB-7 compartments (Figure 5).

Our results support a model where SID-1 is required to facilitate dsRNA transport into the cytoplasm directly from extracellular spaces as well as from endocytotic vesicles. Indeed, other organisms capable of systemic RNAi may be more reliant on endocytosis of dsRNA, including organisms that lack SID-1 (Saleh *et al.* 2006), and at least one insect species has been shown to use both SID-1 and endocytosis (Cappelle *et al.* 2016). A better understanding of the mechanisms by which circulating RNA reaches the next generation will help us identify endogenous inherited RNAs.

## Materials and Methods

### Strains

The following strains were used: N2 wild-type, HC977 *sid-1(qt101)*, HC970 *sid-1(qt78)*, *sid-1(qt78)*; *sid-2(qt40)*, HC306 *sid-2(qt40)*, HC770 *sid-3(tm342)*, HC302 *sid-5(qt24)*, HC975 *sid-1(qt78)*; *sid-3(tm342)*, HC976 *sid-1(qt78)*; *sid-5(qt24)*, DH1390 *rme-2(b1008)*, HC1064 *rme-2(b1008)*; *sid-1(qt101)*, HC17 bIs1 [*vit-2::GFP* + *rol-6(su1006)*]; *emb-27(g48)*, HC1060 *sid-1(qt101)*; bIs1[*vit-2::GFP* + *rol-6(su1006)*], RT123 pwIs21 [*unc119*(+); P*pie-1*::*GFP*::*rab-7*], HC1099 *sid-1(qt78)*; pwIs21[*unc-119*(+); P*pie-1*::*GFP*::*rab7*]

### RNAi

Injections of *unc-22* dsRNA were done at a concentration of approximately ~2 mg/ml. For pseudocoelom injections, the needle was inserted beyond the bend of the gonad arm but before the pharynx or else in the tail beyond the gonad. Injections were done at ~13-20 psi, with successful injections appearing to briefly “highlight” tissues along the entire length of the animal under DIC. For RNAi experiments involving a cross, injected animals were recovered together on a single OP50 plate for ~12 h before the addition of triple the number of appropriate males. After ~36 h of mass mating, individual injected hermaphrodites were singled to new plates along with 3 males and allowed to lay eggs for ~48 h before all P_0_ animals were removed. For the feeding RNAi timing experiment, a mixed population of N2 animals was bleached in a basic sodium hypochlorite solution until adult bodies had dissolved, and the released embryos were rinsed in M9. Embryos were allowed to hatch in shaking M9 for 10 h, and the hatched L1s were then roughly partitioned and transferred to *unc-22* RNAi food (Timmons *et al.* 2001) or OP50 plates as appropriate by pipetting. Each subsequent day, animals were washed off of the plates and washed four times in M9 and then transferred to new appropriate bacteria plates. Gravid adults were washed again, and 30 animals from each group were picked to individual OP50 plates for F_1_ collection. Adult and L4 feeding assays were similarly washed before F_1_ collection, but parents were simply picked from mixed populations. Day 1 adults were prepared by isolating L4 larvae and maintaining for 12 h at 20°C. Scoring for the strong twitching phenotype characteristic of *unc-22* silencing was done in 10 mM levamisole in M9 buffer once the F_1_ progeny were young adults.

### Labeled dsRNA preparation

5EU and Cy5 labeled RNAs were synthesized using the Ampliscribe T7 Flash Transcription Kit (Epicentre) using a modified version of the manufacturer’s protocol to include the substituted nucleotides. Detailed procedures can be found in the supplemental methods.

### Immunohistochemistry

Slides for microscopy were prepared as in (Hinas *et al.* 2012), with some modifications for click labeling of 5EU. For mounted adults, 5EU was conjugated to an Alexa 594- azide using the Click-iT RNA Imaging Kit (Invitrogen). For embryos, 5EU was first conjugated to biotin-azide (Lumiprobe) using the same Click-iT kit, followed by the Alexa Fluor 594 Tyramide SuperBoost Kit with streptavidin (Invitrogen) for enhanced signal. Detailed procedures can be found in the supplemental methods.

### Microscopy

Most images were captured using a Zeiss LSM880 microscope using the ZEN software (Zeiss) at the Harvard Center for Biological Imaging. Embryo images from Cy5/5EU dual labeling experiments were captured using a Zeiss Axiovert 200 spinning disk confocal microscope with Axiovision (Zeiss).

## Supplemental methods

### Labeled RNA preparation

Purified RNA was synthesized by *in vitro* transcription (IVT) from a DNA template containing T7 promoter sequences. For labeled and unlabeled *unc-22* dsRNA, template DNA was prepared by PCR using primers with the T7 promoter sequence appended to the 5’ end of both forward and reverse primers. For the Cy5- and 5EU-containing RNA in Fig. 4, the forward and reverse strands of the eventual dsRNA were prepared independently from template DNA containing only a single T7 promoter.

IVT was performed using an Ampliscribe T7 Flash kit (Epicentre) according to manufacturer instructions with the following modifications for labeled RNAs. For 5EU-labeled RNA, the 100 mM UTP in the reaction was substituted with an equal volume of a 1:1 mixture of UTP and 100mM 5-ethynyl-UTP (Jena Bioscience). 5EU-labeled RNA can also be synthesized using only 5-ethynyl-UTP and no unmodified UTP and behaves similarly in the experiments we performed. For Cy5-labeled RNA, instead of individually adding 0.9 μl of each NTP, a mixture of equal amounts of 100 mM ATP, CTP, and GTP was prepared, and 4.55 μl of this mixture was added to the reaction, followed by 1 μl of 100 mM UTP and 3 μl of 10 mM Cy5-UTP (Amersham). IVT reactions of labeled RNA were incubated at 37°C for 4.5 h, and then treated with DNase I at 37°C for 15 min. Synthesized RNA was purified using an RNeasy kit (Qiagen). Equal quantities of complementary strands of RNA were then annealed together by heating mixtures to 95°C for 2 min and then gradually lowering the temperature to 20°C at a rate of 0.1 °C per second.

### Immunohistochemistry

After injection of 5EU-labeled RNA, adults or isolated embryos were transferred to a poly-L-lysine coated slide and covered with 10 μl of 4% paraformaldehyde solution (PFA). A square cover glass was then placed on to the sample, and excess liquid removed by wicking to a tissue paper until the point of mechanical rupturing of eggshells or cuticles to provide access to the fixative. Samples were then left to fix for 15 min at room temperature before flash freezing in liquid nitrogen. Afterwards, the cover glass was removed with a razor blade to help remove cuticles and eggshells. Samples were then fixed in -20°C methanol for 15 min, and then briefly rinsed in PBS. The final preparation step before labeling was sample permeabilization in a 0.1% Tween-20 in phosphate buffered saline (PBS) solution for 15 min at room temperature.

Labeling of samples containing 5EU-RNA was performed with a Click-iT RNA Imaging kit (Invitrogen) according to manufacturer instructions, but reaction volumes reduced to 100 μl per slide. For adult worm samples, labeling was done with Alexa Fluor 594 azide (Thermo Fisher), but embryo samples typically substituted 2.66 μM biotin azide (Lumiprobe) for additional signal amplification. Click labeling reactions were incubated at room temperature for 30 min and then stopped with addition of stop solution from the kit, and then washed 3 times in PBS. Samples to be labeled only with Alexa fluor could be mounted in Vectashield with DAPI and sealed under a coverslip at this point, or further processed with additional labeling.

Embryo samples requiring signal amplification were further processed with the Alexa Fluor 594 Tyramide SuperBoost Kit with streptavidin (Invitrogen) according to manufacturer instructions, with the tyramide labeling reaction allowed to proceed for 8 minutes. After washing 3 times in PBS for 10 minutes each, slides were mounted and sealed, or additionally labeled by antibody staining.

Samples requiring antibody labeling were incubated at room temperature for 1 h with a rabbit polyclonal GFP antibody (Invitrogen) diluted 1:200 in a solution of 5% bovine serum albumin (BSA) in PBS with 0.1% Tween-20. After washing 3 times in PBS, samples were then incubated at room temperature for 1 h with an Alexa Fluor 488-conjugated goat anti-rabbit secondary antibody (Invitrogen). After washing 3 times in PBS, slides were mounted with Vectashield with DAPI and sealed.

**Figure S1.**
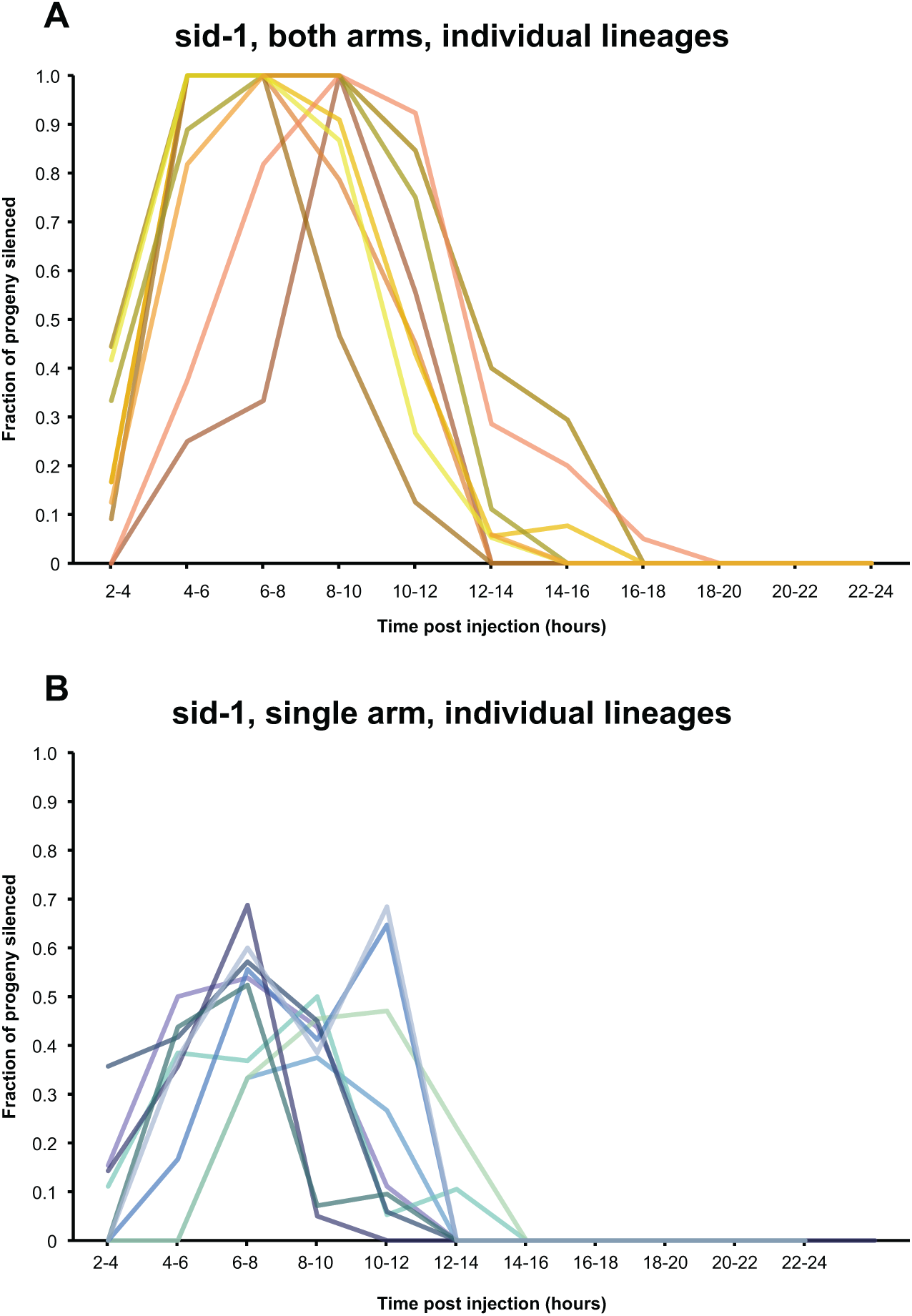
Individual lineages achieve maximum silencing with variable timing. Expanded data from Figure 1, panels 1A and 1C. Each panel in Figure 1 represents the average of two experiments, with the data corresponding to the progeny of each 10 injected animals in each duplicate. Here, the data from one of the replicates from panels 1A and 1C respectively are expanded in (A) and (B) to show the data from each individual injected animal as a separate line.

